# PIEZO1 discriminates mechanical stimuli

**DOI:** 10.1101/2022.06.23.497409

**Authors:** Alper D. Ozkan, Tharaka D. Wijerathne, Tina Gettas, Jérôme J. Lacroix

## Abstract

PIEZO1 channels open in response to numerous mechanical stimuli, such as physical membrane deformations, which modulate the curvature of flexible domains called blades. Yet, whether different stimuli cause similar blade motions and whether these rearrangements correlate with pore opening remain unclear. Here, we scan local conformational changes along the PIEZO1 blade using fluorescent probes. We identify two distant probes, one intracellular proximal and the other extracellular distal, which independently and robustly respond to flow stimuli. Flow-induced signals from both probes exquisitely correlate with PIEZO1-dependent calcium influx and specifically increase in presence of fast-inactivating pore mutations. In contrast, both probes remain fluorimetrically silent to hypotonic shocks and indentations, two stimuli that otherwise evoke normal electrochemical responses in both engineered channels. This study reveals that flow-induced blade motions are functionally coupled to the pore and that at least two distant blade regions discriminate flow from two other stimuli, suggesting that PIEZO1 mobilizes distinct mechanisms to sense a broad range of mechanical cues.

**Teaser:** Fluorimetric evidence suggests that different mechanical stimuli impart distinct rearrangements in PIEZO1’s mechanosensory domains.

## Introduction

Mechanosensitive PIEZO channels couple mechanical forces to intracellular signaling, enabling organisms to control tissue growth, regulate the flow and pressure of internal fluids, and map the topography of their environment (*1*). PIEZO1, one of only two vertebrate PIEZO members (*2, 3*), responds to a bewildering diversity of mechanical stimuli. These include membrane stretch (*4–6*), hydrostatic pressure (*7*), fluid shear stress (*8–13*), intracellular traction forces (*14*), hypotonic shocks (*6, 15, 16*), nano-scale substrate displacements (*17*), mechanical indentations with a microprobe (*2*), and low-intensity ultrasound (*18, 19*). This broad mechanical sensitivity mirrors the breadth of physiological functions governed by PIEZO1 across cells, organs, and physiological systems (*9, 11, 12, 20–30*).

PIEZO1 possesses a homotrimeric structure encompassing a central pore region and three non-coplanar transmembrane blade-like domains, conferring the channel a unique bowl shape (*31–35*). Physical manipulations of the PIEZO1 blade alter channel sensitivity to mechanical forces (*36–39*), suggesting that this domain senses mechanical stimuli by changing its conformation. Indeed, high-speed atomic microscopy kymographs reveal that the PIEZO1 blades reversibly flatten under compression, while cryo-electron microscopy images of PIEZO1 reconstituted in proteoliposomes show that the curvature of its blades changes as a function of the vesicle’s intrinsic curvature (*40, 41*). Molecular dynamics simulations further indicate that stretching (*42*) or flattening (*43*) the lipid bilayer causes PIEZO1 to flatten its blades and open its pore.

Yet, in spite of these efforts, direct experimental evidence for the coupling between PIEZO1 blade motions and pore opening remains scarce. Indeed, although a recent structure captures PIEZO1 with coplanar blades, this structure has a low resolution in the pore, making it difficult to link this flat conformation to a defined functional state (*41*). In addition, the complete loss of mechanosensitivity incurred by covalently crosslinking the blade to an extracellular cap domain above the pore could be caused by a lack of mobility in the cap (*37*). Actuation of the blade with cleverly engineered magnetic nanoparticles modulates current kinetics, but fails to alter open probability in absence of mechanical stimulus (*38*). Genetic deletion of the blade abolishes mechanosensitivity (*36*), yet claims that the blade is sufficient *per se* to confer mechanosensitivity by transplanting this domain to a naïve trimeric channel have been disputed (*44, 45*).

Whether or not opening of the pore is specifically caused by conformational changes in the blade, it remains unclear whether different mechanical stimuli cause similar conformational rearrangements of the channel structure. Indeed, certain blade mutations or deletions differentially alter PIEZOs’ sensitivity to membrane stretch and mechanical indentations (*35, 39, 46*). In addition, PIEZO1 activation by fluid shear stress depends on the presence of intracellular cytoskeletal and/or extracellular matrix filaments (*47, 48*), whereas its activation by membrane stretch does not (*4, 6*).

To address these questions, here we track force-induced rearrangements of the PIEZO1 blades *in cellulo* using site-specific fluorimetry, enabled by the exquisite sensitivity of fluorescent probes to local protein motions. We show that two independent probes, located in the blade ~20 nm apart from each other (according to the AlphaFold2 full-length PIEZO1 structural model (*49*)), produce robust fluorimetric signals in response to fluid shear stress, but not to hypotonic shocks or mechanical indentations, despite the effectiveness of all these stimuli to elicit PIEZO1-dependent electrochemical responses.

## Results

### Generation of PIEZO1-cpGFP constructs

Site-specific fluorimetry, the detection of changes in fluorescence emission from chromophores attached to specific protein sites, has enabled tracking conformational changes in voltage-gated and ligand-gated ion channels (*50, 51*). Here, we adapt this technique to mechanosensitive PIEZO1 channels. To this aim, we used circular permuted green fluorescent proteins (cpGFP) as tractable conformation-sensitive probes. Indeed, conformational changes in the host protein backbone near the site of cpGFP insertion often lead to reduction (quenching) or increase (dequenching) of cpGFP fluorescence emission, through subtle chemical changes in the chromophore vicinity (*52*). Although the degree of cpGFP fluorescence modulation is generally not quantitatively related to the physical displacement of the host protein backbone, these fluorescence changes enable tracking local conformational changes, especially those undergone by sensory domains upon interaction with external stimuli. This property has inspired the development of numerous genetically encodable fluorescent indicators to monitor chemical and physical parameters *in vivo* and with high temporal resolution (*53–58*).

We cloned cpGFP from the voltage-indicator ASAP1 (*59*) and inserted it at the carboxyl end of mouse PIEZO1 (mPIEZO1) residues 86, 300 and 1591. These positions are spread along the long blade, which is divided into 9 motifs called PIEZO repeats A-I (**Figure 1A**), three positions known to tolerate proteinogenic modifications with no major functional impacts (*4, 15, 38*). We also inserted cpGFP at residue 656, i.e. adjacent to an extracellular loop necessary for mechanical activation (*35*); and at residue 1299, i.e. at the distal end of a long intracellular beam anticipated to transmit force from the blade to the pore (*35, 39*).

**Figure 1.**
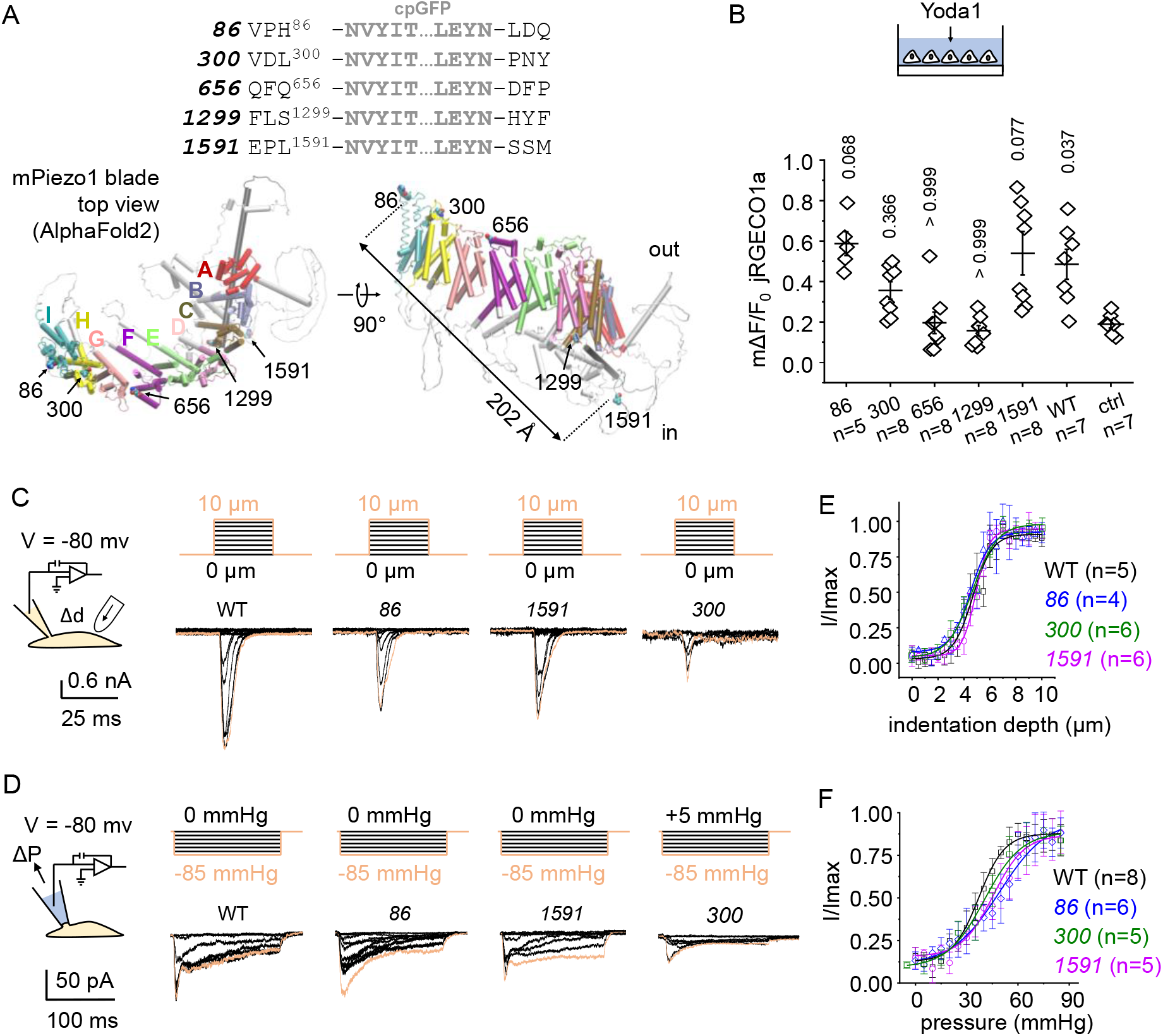
Functional characterization of cpGFP constructs. (**A**) *Top*: amino acid sequences showing cpGFP insertion sites into mPIEZO1. *Bottom*: position of cpGFP insertion sites in the full-length AlphaFold2 structural model of the mPIEZO1 blade (letters indicate PIEZO repeats) (*49*). The figure also indicates the Cα-Cα distance between H86 and L1591. (**B**) Maximal calcium response (mΔF/F0) to acute perfusion with 100 μM Yoda1 from HEK293T^ΔPZ1^ cells transfected with the jRGECO1a plasmid only (ctrl) or co-transfected with plasmids encoded ethe indicated constructs. Dots correspond to the mean mΔF/F0 value obtained from at least 20 cells from independent wells. Numbers above bars indicate p-values from Kruskal-Wallis test with Dunn’s multiple comparisons. (**C-D**) Representative macroscopic current traces of *86*, *300*, *1591*, and WT mPIEZO1 in response to poking (C, indentation depth: 0 to 10 μm) or mechanical stretch (D, pipette pressure: +5 to −85 mmHg). (**E-F**) Relative peak current amplitude (I/Imax) plotted as a function of poking displacement (E) or pipette pressure (F) in indicated constructs. In panels E and F, lines are Boltzmann fits to the data. In panels B, E, F, error bars = s.e.m.

We first tested whether the presence of the cpGFP at these positions impact channel function by measuring the sensitivity of our constructs to the chemical agonist Yoda1 and to two mechanical stimuli. To this aim, plasmids encoded our constructs (named *86*, *300*, *656*, *1299* and *1591*) were individually transfected into mechano-insensitive HEK293T^ΔPZ1^ cells in which endogenous human PIEZO1 expression is abolished (*60*). Sensitivity to 100 μM Yoda1 was assessed using a standard calcium imaging assay (*61*) by co-transfecting cells with a plasmid encoding the red calcium indicator jRGECO1a to avoid spectral overlap with cpGFP (*62*). The maximal calcium-dependent relative fluorescence change, or mΔF/F0, measured in cells transfected with the jREGECO1a plasmid only was similar to the mΔF/F0 measured in cells co-transfected with *656* or *1299* plasmids (Kruskal-Wallis test with Dunn’s test for multiple comparisons p-values > 0.99) (**Figure 1B**), suggesting these two constructs are not functional. In contrast, mΔF/F0 was higher in cells co-transfected with a plasmid encoding either of the other constructs, an effect statistically significant for WT, *86*, and *1591* (0.037 < non-parametric p-values < 0.077).

We next used cell-attached pressure-clamp and whole-cell poking to mechanically evoke ionic currents in constructs *86*, *300*, and *1591* (*63*) (**Figure 1C–D**). Because PIEZO currents inactivate (*3*), we used the relative peak of mechanically-induced current (I/Imax) as a proxy for mechanosensitivity. The curve plotting I/Imax as function of pipette pressure or indentation depth are visually very similar between the three engineered constructs and unmodified wild-type (WT) mPIEZO1 channels (**Figure 1E-F**). Although construct *300* exhibits normal mechanosensitivity, it yields smaller peak currents relative to the others, mirroring its weaker sensitivity to Yoda1 (**Figure 1B**). We thus mainly focus our efforts on characterizing *86* and *1591*.

### *86* and *1591* are fluorimetrically sensitive to flow

We seeded transfected cells into laminar flow chambers to test the sensitivity of our constructs to shear stress, which we delivered by perfusing the chamber with Hanks’ Balanced Salt Solution (HBSS) at a calibrated flow rate (see Materials and Methods). A series of intermittent flow pulses (10 s on / 10 s off) of incrementally increasing amplitude (from 0.02 to 0.69 Pa) elicits robust and transient dequenching of cpGFP fluorescence in cells expressing either *86* or *1591* (**Figure 2A and Movies S1-2**). We previously showed that the same flow protocol fails to elicit fluorescence signals in cells expressing the cpGFP-carrying proteins ASAP1 or Lck-cpGFP, a control protein that consists of a fusion between cpGFP and the membrane bound N-terminal domain of the Lck kinase (*53*). As example, we show representative traces obtained with cells expressing Lck-cpGFP (**Figure 2A**). The inability of flow stimuli to elicit cpGFP signals in these two independent control constructs indicate that the signals produced by *86* and *1591* depend on PIEZO1 acting as the host protein for cpGFP. This means that these signals are unlikely to be caused independently of conformational changes in PIEZO1, for example as a result of intermolecular collisions between cpGFP and surrounding lipids.

**Figure 2.**
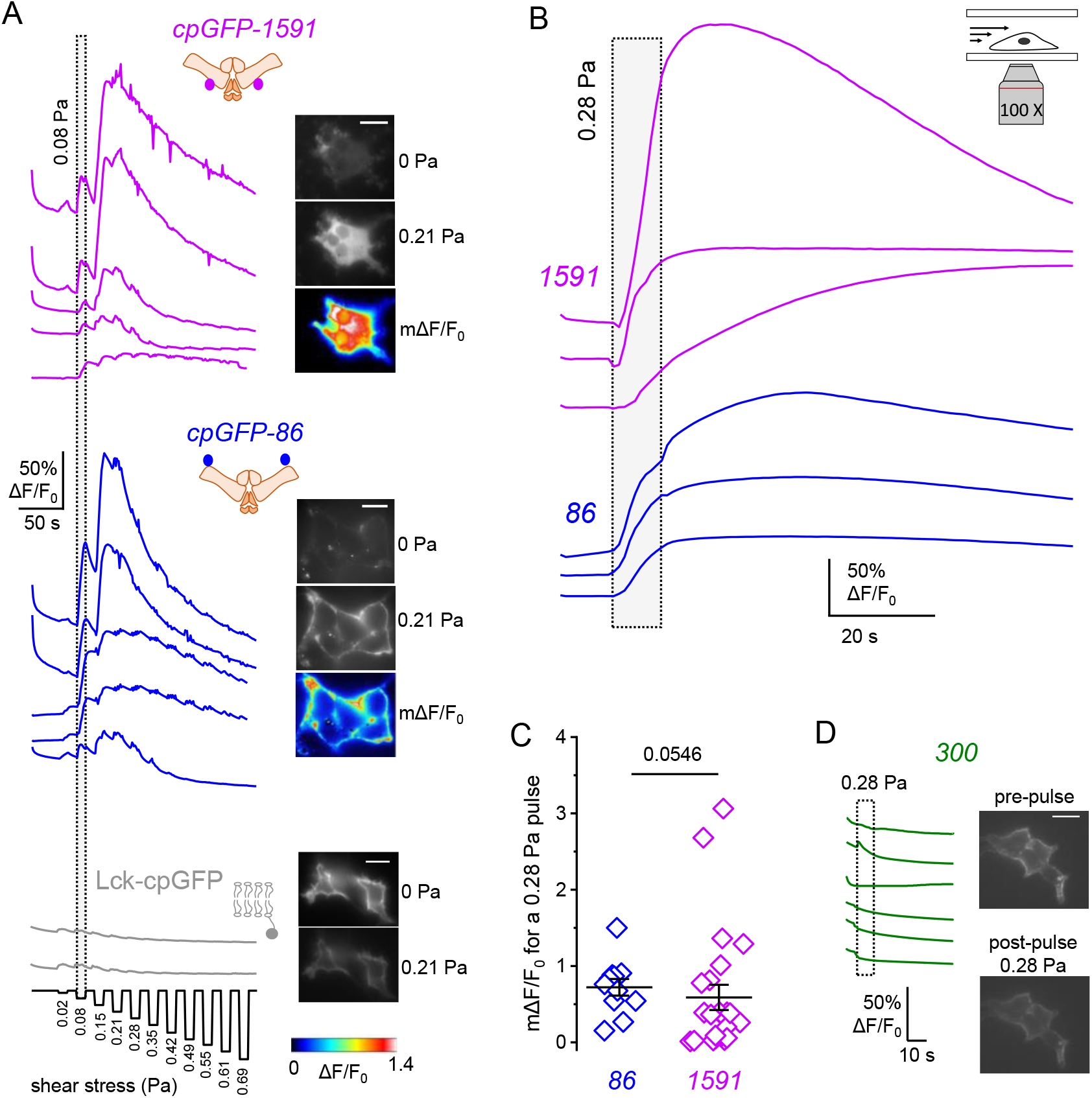
Flow stimuli induce robust fluorescent responses in both *86* **&***1591*. (**A**) Representative cpGFP fluorescence traces, static snapshots and mΔF/F0 images from cells expressing *86*, *1591*, or Lck-cpGFP, in response to shear stress applied in a series of 11 escalating flow pulses (duration: 10 s, magnitude: 0.022 – 0.691 Pa). (**B**) Representative cpGFP fluorescence traces obtained from cells expressing *86* or *1591* and stimulated with a 10 s / 0.28 Pa pulse. (**C**) Bar graphs representing mΔF/F0 values obtained from cells expressing *86* or *1591* using experiments shown in (B). The p-value is obtained from a Mann-Whitney U-test and the errors bars are s.e.m. (**D**) Representative cpGFP fluorescence traces from cells expressing *300* and stimulated with a 10 s / 0.28 Pa pulse. In panels A and D, scale bars = 10 μm. All displayed data were obtained from independent flow chambers.

We next used individual flow pulses to evoke these fluorimetric responses. Single pulses of 10 s duration and 0.28 Pa amplitude produce large dequenching signals from both probes (**Figure 2B**). The time course and amplitude of these signals vary from cell to cell. This heterogeneity is not surprising, as transfected cells visually exhibit variable morphology, polarity and adhesion profile, factors anticipated to affect the mechanical stress experienced by fluid shear stress (*64*). Nevertheless, the mean maximal amplitude of these signals were similar between *86* (+75 ± 11 %) and *1591* (+59 ± 17 %) (p-value = 0.0546) (**Figure 2B-C**). Using the same stimulus, no such signals were detected in cells expressing *300*, although these cells display membrane cpGFP fluorescence, suggesting proper folding of the chromophore and proper targeting at the cell membrane (**Figure 2D**), as anticipated from electrophysiology data (**Figure 1C-F**). The fact that only two out of three constructs respond to flow demonstrates that the property of the cpGFP probe to light up in response to flow strictly depends on its position within the blade, as expected if these signals were precisely produced by local rearrangements in this domain.

### Flow-induced fluorimetric signals correlate with functional channel transitions

Calcium imaging experiments show that our 10 s / 0.28 Pa flow pulse elicits larger calcium signals in cells co-transfected with plasmids encoded jRGECO1a and WT mPIEZO1, *86*, or *1591* compared to cells transfected with the jRGECO1a plasmid only (0.0231 < non-parametric p-values < 0.0387) (**Figure 3A-C**), showing that this standard flow stimulus is effective to activate PIEZO1. We next used dual-wavelength imaging to show that both conformation-dependent cpGFP signals and calcium-dependent jRGECO1a signals occur together in the same cells (**Figure 3D**). Indeed, the time-course of cpGFP and jRGECO1a fluorescence are highly positively correlated during the duration of the flow pulse, evidenced by Pearson’s correlation coefficients values near unity (0.86 ± 0.06, n = 8, for *86* and 0.97 ± 0.01, n = 6, for *1591*) (**Figure 3E-F**). The two fluorescence signals appear sometimes negatively correlated before or after exposure of the cells to flow, showing that the strong positive correlation observed during the pulse cannot be primarily caused artefactually, for instance by photon leakage across the spectrally separated optical paths collecting light from cpGFP and jRGECO1a. Taken together, these results show that, at our experimental temporal resolution of 1 frame s^−1^, cpGFP signals occur concomitantly with PIEZO1-dependent calcium influx, as expected if the cpGFP signals correlate with flow-induced conformational changes leading to opening of the pore.

**Figure 3.**
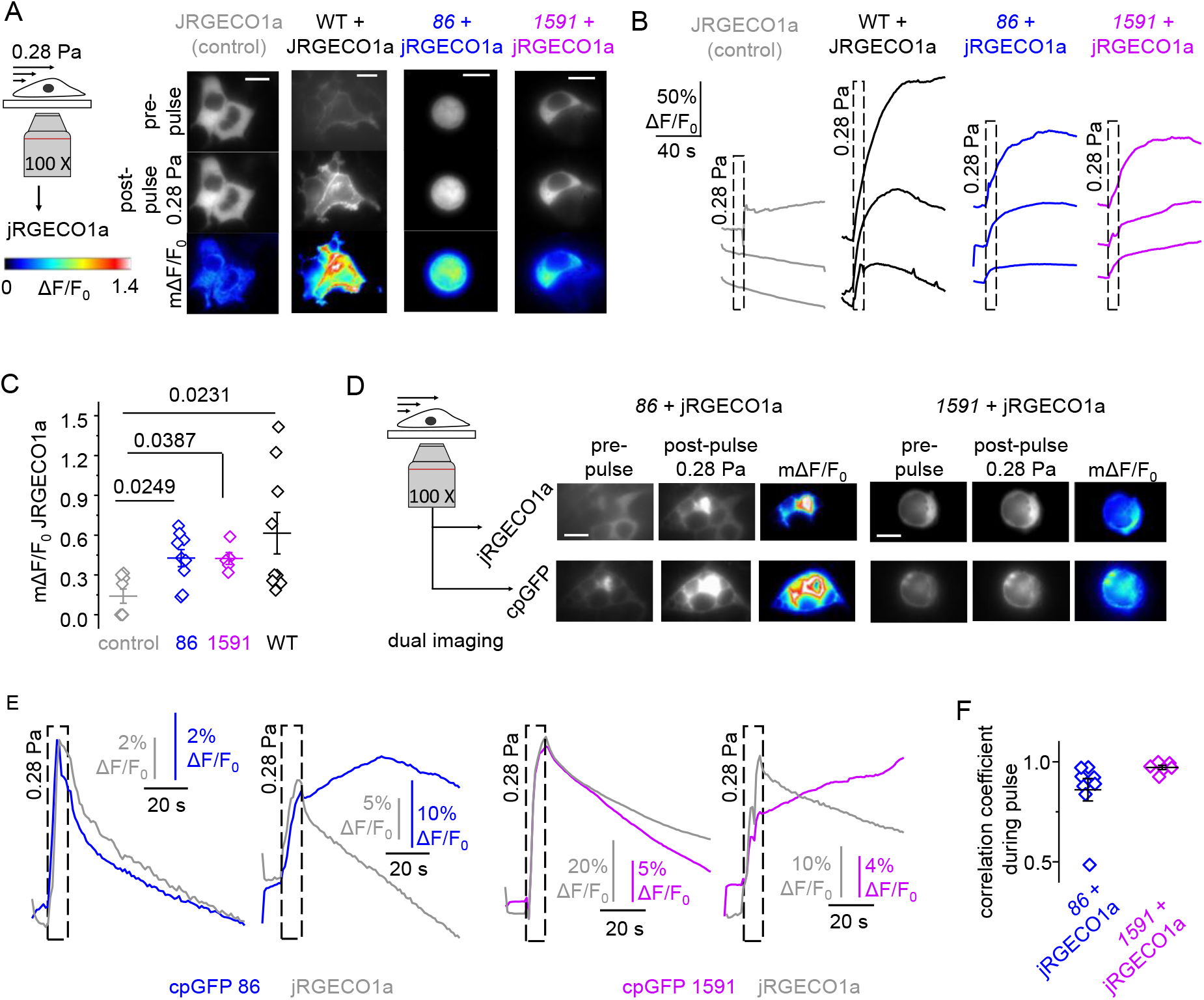
Flow-induced cpGFP signals in both 86 & 1591 correlate with calcium uptake. (**A-B**) Representative images (A) and time traces (B) of jRGECO1a fluorescence in cells transfected with jRGECO1a and co-transfected or not (control) with *86*, *1591*, or WT mPIEZO1 and exposed to a 10 s / 0.28 Pa flow pulse. (**C**) Bar graphs representing WT-normalized jRGECO1a mΔF/F0 values for indicated co-transfection conditions from experiments described in (A-B). Numbers above bars indicate p-values from Kruskal-Wallis test with Dunn’s multiple comparisons. (**D**) Dual fluorescence (jRGECO1a/cpGFP) images and time traces showing the concurrence of cpGFP dequenching and cytosolic calcium entry in cells transfected with *86* or *1591*. Scale bars = 10 μm.

To further determine a link between cpGFP signal and channel function, we introduced pairs of mutations in the pore region that are known to slow down or accelerate the rate of inactivation of macroscopic currents (Tau_inactivation_). To slow down Tau_inactivation_, we introduced the mutation pair M2241R and R2482H (MR/RH), murine homologs of human mutations M2225R (located in the cap) and R2456H (located in the inner pore helix) which both slow down inactivation (*65, 66*). The second pair consists of mPIEZO1 mutations L2475I and V2476I (LI/VI) (both located in the inner pore helix) which individually accelerate inactivation (*67*). Because Tau_inactivation_ varies as a function of the membrane potential (*3*), we measured Tau_inactivation_ at −80 mV and +80 mV in these mutants and in their unmodified parent cpGFP constructs using whole-cell poking recordings. Introducing MR/RH in both *86* and *1591* slows down Tau_inactivation_ by ~1.5-fold at +80 mV and by ~10-fold at −80 mV relative to their respective parent constructs (**Figure 4A-B**), whereas inserting LI/VI in both *86* and *1591* accelerates Tau_inactivation_ by ~10-fold at +80 mV and by ~2-fold at −80 mV relative to their respective parent constructs (**Figure 4C-E**). Remarkably, the maximal amplitude of cpGFP signals induced by a 10 s / 0.28 Pa pulse increases in the presence of fast-inactivating LI/VI mutations relative to unmodified controls for both constructs, an effect that was much larger for *86* (+280 ± 35 % vs. +33 ± 6 % for control, p-value <0.0001) than for *1591* (+112 ± 15 % vs. 32 ± 11 % for control, p-value = 0.020). This increase of signal amplitude seems to correlate with the development of a slow kinetic component of the cpGFP fluorescence signal, which is absent in unmodified constructs or in constructs harboring the MR/RH mutations (**Figure 4D**). The origin of the LI/VI mutant phenotype is unclear. Analysis of electrophysiology data suggests that this phenotype is likely not due to an increase of protein expression, as current densities were similar in all tested constructs (p-values > 0.479) (**Figure 4F**).

**Figure 4.**
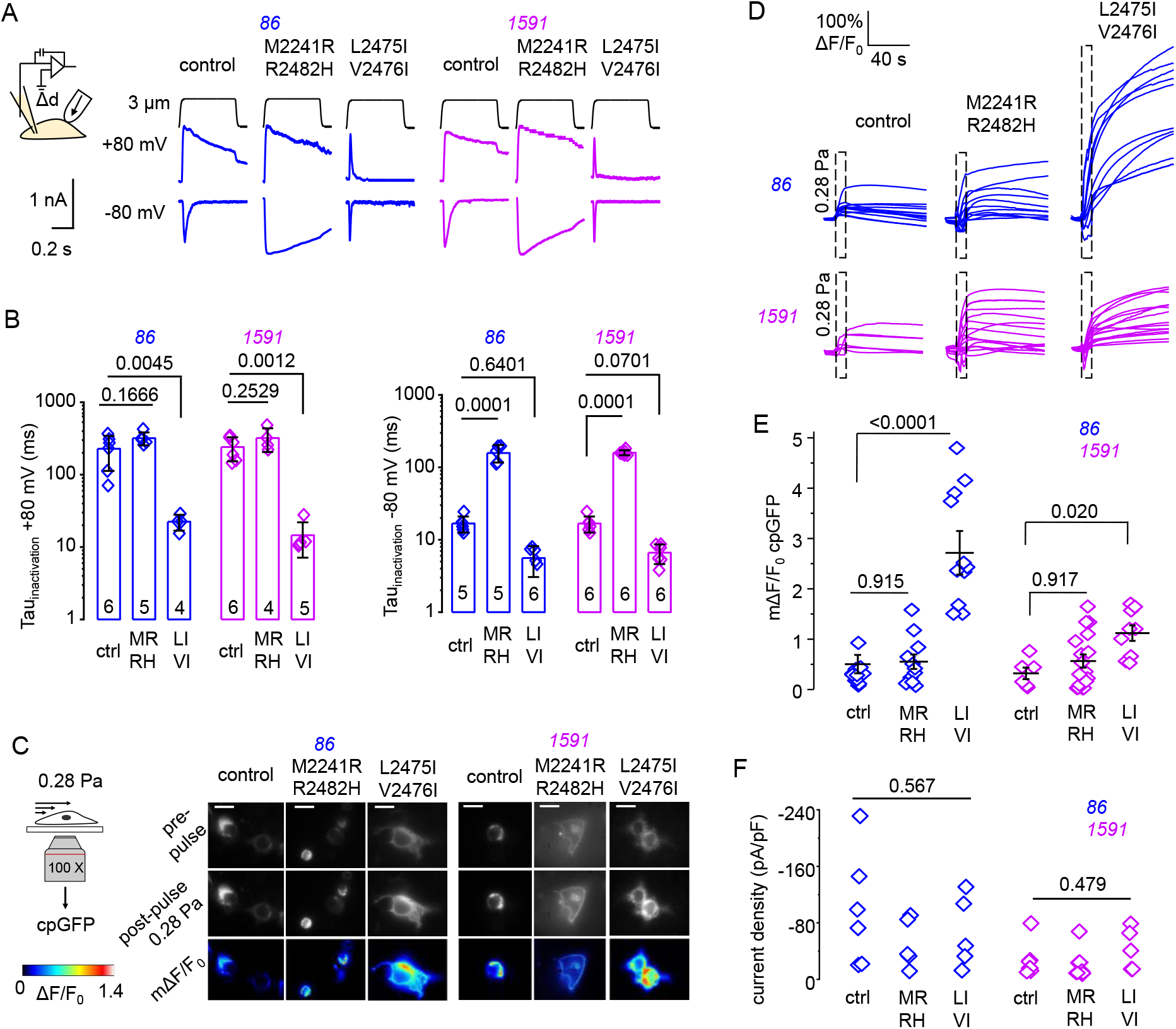
Modulation of cpGFP signals by fast-inactivating mutations. (**A**) Representative macroscopic poking-evoked current traces from cells expressing *86* or *1591*, modified or not (control) with pairs of mutations predicted to increase (L2475I/V2476I) or decrease (M2241R/R2482H) Tau_inactivation_. Cells were stimulated using 3 μm poking stimuli for 500 ms at a holding potential of −80 mV or +80 mV. (**B**) Tau_inactivation_ obtained using experiments described in (A) plotted for unmodified *86* and *1591* constructs (Ctrl) and for *86* and *1591* constructs carrying mutations (MR/RH = M2241R/R2482H; LI/VI = L2475I/V2476I). (**C**) Representative epifluorescence images of cells expressing modified and unmodified *86* and *1591* constructs before and after stimulation with a single 10 s / 0.28 Pa flow pulse. Scale bars = 10 μm. (**D**) Time course of cpGFP fluorescence traces obtained from cells transfected with indicated constructs and stimulated with a single 10 s / 0.28 Pa pulse. (**E**) Scatter plot showing mΔF/F0 values from experiments described in (D). (**E**) Current density scatter plots for data obtained in (A). Each point in box plots represents data from independent experiments. Numbers above plots in panels (B), (E), and (F) indicate p-values from Kruskal-Wallis tests with Dunn’s multiple comparisons.

### cpGFP probes are fluorimetrically insensitive to hypotonic shocks and indentations

We next tested whether our probes fluorimetrically respond to other stimuli. We first acutely exposed transfected cells to a hypotonic shock of ~58 mOsmol L^−1^. This osmotic stimulus produced rapid PIEZO1-dependent calcium signals in cells co-transfected with jRGECO1a and with WT mPIEZO1, *86*, or *1591* compared to cells transfected with jRGECO1a alone (non-parametric p-values < 0.0006) (**Figure 5A-B**). However, this stimulus was ineffective at eliciting detectable cpGFP dequenching in *86*, *300*, or *1591*, as cpGFP fluorescence emission in each construct tends to monotonically decay before and during hypotonic stimulation, indicative of photobleaching effects (**Figure 5C**). The lack of cpGFP response occurs despite osmotic swelling of the cells, which is clearly noticeable from visual examination of imaging data (**Movies S3-5**).

**Figure 5.**
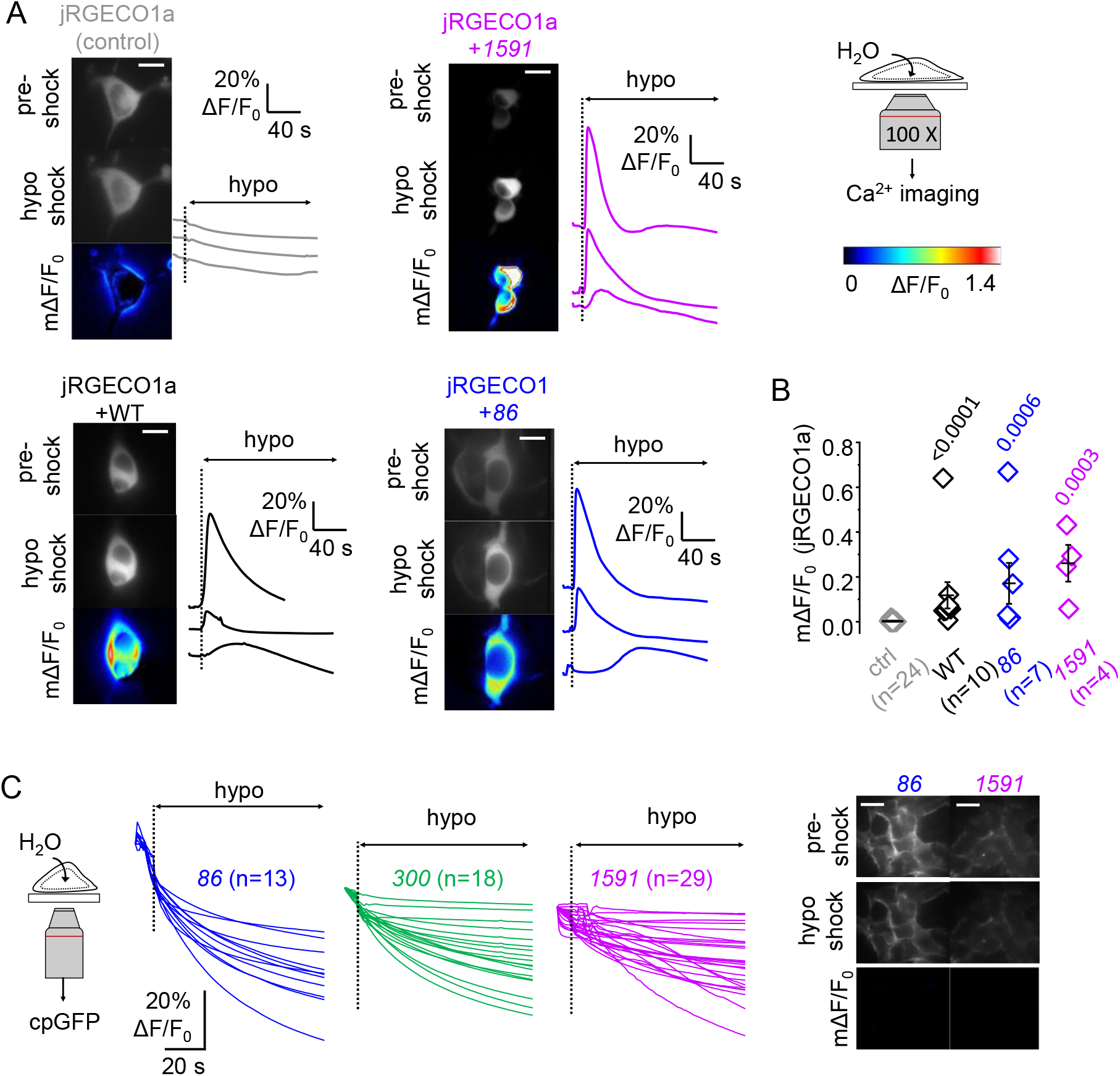
*86* & *1591* are fluorimetrically silent to hypotonic shocks. (**A**) Representative jRGECO1a fluorescence images and time traces in cells transfected with a jRGECO1a plasmid only (control) or co-transfected with *86*, *1591*, or WT PIEZO1, and exposed to a hypotonic solution (~58 mOsmol L^−1^). (**B**) Scatter plots showing jRGECO1a mΔF/F0 values from experiments described in (A). Each point in box plots represents a cell or group of cells from independent experiments. Numbers above plots in indicate p-values from Kruskal-Wallis test with Dunn’s multiple comparisons. (**C**) Representative cpGFP fluorescence images and time traces of *86*, *300*, and *1591* following exposure to a hypotonic solution. In panels (A) and (C), scale bars = 10 μm.

We next tested whether our probes fluorimetrically respond to cellular indentations. This stimulus evoke normal ionic currents with a saturation of I/Imax plots at approximately 7 μm probe displacement (**Figure 1**). As expected, poking stimuli of 7 μm displacement and 1500 ms duration evoke larger calcium signals in cells co-transfected with jRGECO1a and WT mPIEZO1, *86*, or *1591*, compared to control cells transfected with jRGECO1a alone (0.0007 < p-values < 0.0890) (**Figure 6A-B**). Interestingly, the calcium-dependent fluorescence signals tend to specifically occur at the site of impact, as expected if these are induced by the mechanical stimulus. However, this stimulus was totally ineffective at evoking cpGFP dequenching in either *86*, *300*, or *1591* (**Figure 6C**). Small increase or decrease of cpGFP fluorescence (mΔF/F0 < ± 10 %) occasionally occur during indentation of cells transfected with these constructs. However, visual examination of imaging data suggests that these effects are due to unpreventable movements of the cell membrane during indentation (**Movies S6-8**). Deeper indentations of 10 μm were still ineffective at evoking fluorimetric cpGFP responses, although they occasionally ruptured the cell membrane (data not shown).

**Figure 6.**
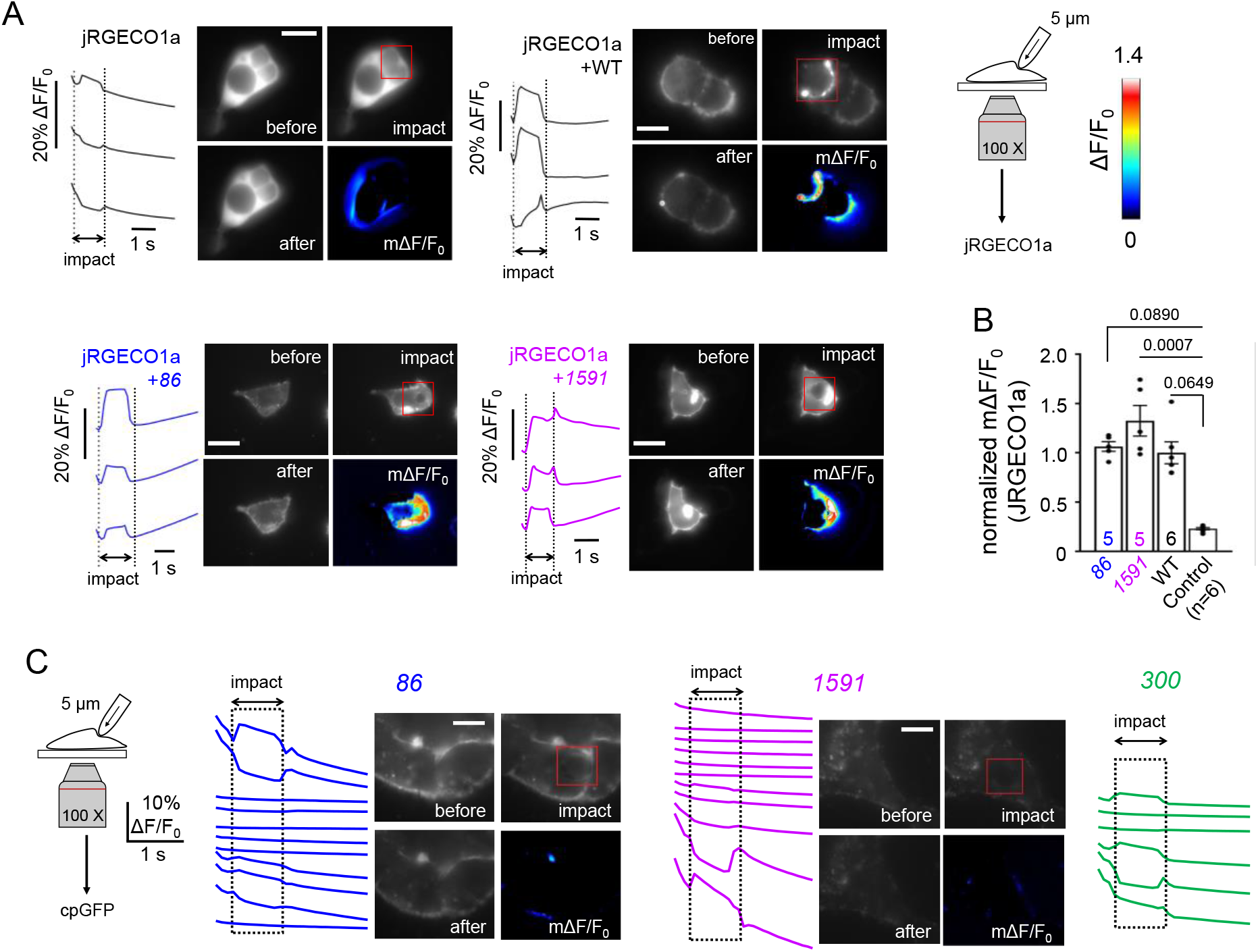
*86* & *1591* are fluorimetrically silent to indentations. (**A**) Representative calcium sensitive epifluorescence images and time-course from cells transfected with a jRGECO1a plasmid and co-transfected or not (control) with *86*, *1591*, or WT mPIEZO1 plasmids and stimulated with a 7 μm poke stimulus (red squares). (**B**) Scatter box plots showing mΔF/F0 values from experiments described in (A) and normalized to mean WT value (error bars = s.e.m.). Each dot represents a cell or group of cells from independent experiments. Numbers above bars indicate p-values from Kruskal-Wallis test with Dunn’s multiple comparisons. (**C**) Representative cpGFP epifluorescence images and time-course from cells expressing *86*, *300*, and *1591* and stimulated with a 7 μm poke stimulus (red squares). In panels (A) and (C), scale bars = 10 μm.

## Discussion

In this study, we provide direct fluorimetric evidence that the PIEZO1 blade acts as a mechanosensory domain, and that this domain discriminates flow from two other mechanical stimuli. This conclusion is based on several independent observations showing that cpGFP signals correlate with conformational changes associated with functional channel transitions. First, flow-induced cpGFP signals occur at two independent blade positions and depend on the probe position within the blade. Second, these signals do not occur in two non-mechanosensitive membrane proteins harboring a cpGFP near the extracellular (as in *86*) or intracellular (as in *1591*) side of the cell membrane (*53*). Third, these signals are quantitatively correlated with pore opening, assessed with calcium imaging. Fourth, these signals are specifically enhanced by pore mutations accelerating inactivation kinetics.

Regarding the fourth observation, although fast-inactivating mutations increase cpGFP signal amplitude, an effect much larger for the distal probe, mutations slowing down inactivation did not produce the opposite effect (a decrease in signal amplitude), making it difficult to associate cpGFP signals with specific conformational or functional transition(s). Regardless, the fact that point mutations in the pore cause a large effect on the fluorescence signal produced by the distal cpGFP probe demonstrates the existence of a long distance (> 20 nm) conformational coupling between the proximal C-terminal pore region and the distal N-terminal tip of the blade, consistent with the blade acting as a mechanosensory domain controlling the opening of activation gates in the pore.

A limitation of site-specific fluorimetry is the impossibility to quantitatively correlate fluorimetric signals with specific protein motions. Corollarily, an absence of fluorimetric signal is difficult to interpret. Indeed, since a limited types of protein rearrangements are predicted to be effective at modulating cpGFP fluorescence (*52*), it is generally not possible to know whether the protein region near the site of cpGFP insertion remains static or not when the probe fails to fluorimetrically respond, such as when our constructs *86* and *1591* are stimulated with osmotic shocks or indentations. It is thus possible that flow prompt distinct blade rearrangements at these positions, motions that happen to fluorimetrically modulate our probes. It is also possible that osmotic swelling and indentations do not induce protein motions at these two regions, explaining the absence of cpGFP signal.

Currently, PIEZO activation by changes in membrane tension and curvature is best explained by a flattening motion of the blades (*40, 41, 43, 68*). Yet, in many cases, experimentally delivered mechanical stimuli (such as those used in this study) are complex in nature and may apply forces possessing tensional, compressional, and/or frictional components, each of them being potentially applied along independent spatial directions relative to the cell membrane. It is thus difficult to reconcile the physical complexity characteristic of experimental mechanical stimuli with the proposition that that all of them act by producing identical changes in membrane tension or curvature.

In line with previous studies showing that loss-of-function mutant phenotypes depend on the nature of the mechanical stimulus (*39, 46*), our work strongly suggests that a unique force-sensing mechanism is unlikely to describe PIEZO1 activation by all its known stimuli. Since PIEZO1’s flow-sensitivity critically depends on the integrity of the cytoskeleton and/or extracellular matrix elements (*47, 48, 69*), flow stimuli may act *via* a force-from filament mechanism, perhaps directly mediated by protein-protein interactions involving intracellular channel domains (*46, 70, 71*). Our observation that blade motions depend on the nature of the mechanical stimulus could thus be explained by the existence of multiple, non-exclusive echanosensory mechanisms, wherein external forces are transmitted to the channel from lipids and/or from filaments.

## Methods

### Molecular cloning

A pCDNA3.1-mPIEZO1 plasmid was generously donated by Dr. Ardèm Patapoutian (Scripps Research). cpGFP fragments were PCR-amplified from a pCDNA3.1-ASAP1 plasmid gifted by Dr. Francois Saint-Pierre (Baylor College of Medicine & Rice University) and inserted to desired positions into the pCDNA.3.1-mPIEZO1 plasmid using High-Fidelity DNA Assembly (New England Biolabs). The presence of cpGFP inserts was confirmed by Sanger sequencing (GENEWIZ). The pCDNA3.1-jRGECO1a plasmid was obtained from a previous study (*53*). The double mutants M2241R-R2482H and L2475I-V2476I were inserted into pCDNA3.1-mPIEZO1-cpGF*86* and pCDNA3.1-mPIEZO1-cpGF1591 using High-Fidelity DNA Assembly and verified by Sanger sequencing.

### Cell culture and transfection

HEK293T^ΔPZ1^ cells were a gift from Dr. Patapoutian. Cells were cultured in standard conditions (37 °C, 5 % CO_2_) in a Dulbecco’s Modified Eagle’s Medium supplemented with Penicillin (100 U mL^−1^), streptomycin (0.1 mg mL^−1^), 10 % sterile Fetal Bovine Serum, 1X Minimum Essential Medium non-essential amino-acids and without L-glutamine. All cell culture products were purchased from Sigma-Aldrich. Plasmids were transfected in cells (passage number < 35) seeded in 96-well plates at ~50 % confluence 2-4 days before the experiment with FuGene6 (Promega) or Lipofectamine 2000/3000 following the manufacturer’s instructions. 1-2 days before experiments, cells were gently detached by 5 min incubation with Phosphate Buffer Saline and re-seeded onto 18 mm round glass coverslips (Warner Instruments) coated with Matrigel (Corning) or onto single or six-channels microfluidic devices (Ibidi μ-slides VI 0.4 or μ-slides I 0.4).

### Fluorescence imaging

Cell culture medium was replaced with HBSS approximately 20 min before imaging experiments. Excitation light was generated by a Light Emitting Diode light engine (Spectra X, Lumencor), cleaned through individual single-band excitation filters (Semrock) and sent to the illumination port of an inverted fluorescence microscope (IX73, Olympus) by a liquid guide light. Excitation light was reflected onto the back focal plane of a plan super apochromatic 100X oil-immersion objective with 1.4 numerical aperture (Olympus) using a triple-band dichroic mirror (FF403/497/574, Semrock). Fluorescence emission from emerald-color transfected cells was filtered through a triple-band emission filter (FF01-433/517/613, Semrock) and sent through beam-splitting optics (W-View Gemini, Hamamatsu). Split and unsplit fluorescence images were collected by a sCMOS (Zyla 4.2,) or by an emCCD (iXon Ultra 897) ANDOR camera (Oxford Instruments). Spectral separation by the Gemini was done using flat imaging dichroic mirrors and emission filters (Semrock). Images were collected by the Solis software (Oxford Instruments) at a rate of 1 frame s^−1^ or 10 frames s^−1^ (for poking experiments). Image acquisition and sample illumination were synchronized using TTL triggers digitally generated by the Clampex software (Molecular Devices). To reduce photobleaching, samples were pulse-illuminated 200 ms per frame during acquisition.

### Image analysis

The first frame of each image stack was initially pre-processed in ImageJ by manually drawing individual cell boundaries and cropping out all background pixels. This mask was then used to define each cell as a unique region of interest. An in-house MATLAB script was then used to determine the average intensity of all pixels associated with each cell, F, and determine mΔF/F0 value for each cell across the trajectory. The MATLAB script is available to download at https://github.com/LacroixLaboratory

### Cell indentation

Premium standard wall borosilicate capillaries 1.5mm × 4 in (Warner Instruments) were heat-pulled on a horizontal puller (Sutter P-97) and fire-polished on a microforge (Narishige MF-900) to produce smooth and round poking probes with tip diameter 2-5 μm. Poking probes were directly mounted onto a closed-loop piezoelectric actuator (LVPZT, Physik Instrumente) attached to a micromanipulator (MP-225, Sutter Instruments). Probes were moved as close to the surface of the cell as possible at an angle of approximately 60° and without physical contact. This initial position corresponds to a 0 μm displacement. Probes were linearly displaced in their longitudinal axis at a speed of 1 μm ms^−1^ using a LVPZT amplifier (E-625.SR, Physik Instrumente) and external voltage triggers commanded by Clampex. For imaging experiments, consecutive pokes were delivered at a rate of one poke per minute.

### Hypotonic shocks

During image acquisition, the extracellular recording solution (HBSS) from cultured dishes was replaced with a hypotonic solution containing 5 mM NaCl, 5 mM KCl, 2 mM MgCl_2_, 1 mM CaCl_2_, 10 mM HEPES (pH 7.4 with HCl or NaOH) and 10 mM Glucose. The osmolarity of the hypotonic solution (~58 mOsmol L^−1^) was measured by a micro-sample osmometer (Advanced Instruments Fiske 210).

### Electrophysiology

Cells were used 2-3 days after transfection. Pipettes were pulled from thin-wall borosilicate capillaries with internal filament (GC 150TF-7.5, Harvard Apparatus) to a resistance of 2-3 MΩ using a vertical pipette puller PC-10 (Narishige). Pipettes were filled with an internal solution containing 140 mM KCl, 10 HEPES, 10 mM TEA and 2 mM EGTA (pH 7.4 with NaOH), whereas HBSS (GIBCO) was used as bath solution. Transfected cells were placed onto an inverted microscope (Olympus IX73) mounted on an air table (TMC). Experiments were performed at room temperature using an Axopatch 200B capacitor-feedback patch clamp amplifier (Molecular Devices) connected to a Digidata 1550B low-noise data acquisition system plus hum silencer (Molecular Devices) and controlled using the Clampex software (Molecular Devices). For cell-attached pressure-clamp recordings, negative pressure stimuli were delivered to the backside of patch pipettes using a Clampex-controlled high-speed pressure clamp (ALA Scientific Instruments, USA). Resting membrane potential of the transfected cells was −28 ± 5 mV when measured using the above-mentioned saline solutions. Therefore, a +50 / −110 mV holding potential was applied through the pipette electrode to provide ~ −80 / +80 mV potential across the membrane patch. For whole-cell poking recordings, pipettes were fire polished using a microforge (MF2, Narishige) and indentation stimuli were delivered as described above. Currents were acquired using a sampling rate of 500 kHz and a filtering frequency of 5 kHz. Data were subsequently filtered at 500 Hz for display.

### Fluid shear stress stimulation and calculations

Fluid shear-stress stimulation was done by circulating extracellular solutions at various flow rate into 0.4 mm height μ-slide flow chambers (Ibidi) using a peristaltic (Golander) or a syringe (NE-1000, New Era Pump Systems) pump TTL-controlled by Clampex. The average amplitude of wall shear stress *τ* applied at the cell surface was estimated using Ibidi’s empirical equation relating τ (Pa) with flow rate Φ (mL min^−1^) and dynamic viscosity *η* (Pa s):

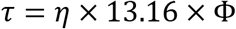

A *η* value of 0.001 Pa s was used for HBSS at room temperature.

### Statistical analyses

We used non-parametric statistical tests to compare experimental groups. The exact nature of each test is indicated in figure legends. All statistical analyses were performed on GraphPad Prism 9.0. For fluorimetry experiments, sample size values (n) indicate single cells or clusters of cells. For electrophysiology, n represents the number of individually patches. All error bars are standard errors of the mean.

## Acknowledgements

This work was supported by National Institute of Health grant GM130834 to J.J.L.

## Author contributions

J.J.L. conceived the project; A.D.O., T.W., T.G, and J.J.L. performed experiments. A.D.O., T.W. and J.J.L. analyzed data; J.J.L. wrote the manuscript.

## Data availability

All data is available in the manuscript and in a Source Data File, deposited online at Open Science Framework (DOI 10.17605/OSF.IO/94F8M).

